# Neural entrainment induced by periodic audiovisual stimulation: A large-sample EEG study

**DOI:** 10.1101/2023.10.25.563865

**Authors:** Joel Frohlich, Ninette Simonian, Grant Hanada, Christian Kothe, Nicco Reggente

## Abstract

Stroboscopic or “flicker” stimulation is a form of periodic visual stimulation that induces geometric hallucinations through closed eyelids. While the visual effects of this form of sensory stimulation have received considerable attention, few studies have investigated the neural entrainment effects of periodic visual stimulation. Here, we introduce two variants of the classic flicker paradigm while recording EEG to study neural entrainment effects in a large sample (over 80 participants per condition). In the first condition, we used multimodal stimulation composed of two simultaneous visual strobe frequencies paired with binaural beats which provided auditory stimulation at roughly the same frequency as the slower strobe. We compared this condition to sham stimulation, in which both strobes were set to very low frequencies and in which the binaural beats were absent. Additionally, we compared both conditions to a control group in which participants focused on their breathing during eyes-closed meditation (no stimulation). Our results demonstrate powerful evidence of neural entrainment at the frequency of the slower strobe in the experimental condition. Moreover, our findings resemble effects reported in prior literature using conventional non-invasive techniques for electromagnetic brain stimulation. We argue that stroboscopic stimulation should be further developed along these lines, e.g., as a potential therapeutic technique in psychiatric disorders.

## Introduction

Noninvasive brain stimulation is generally delivered using forms of energy that are not directly detectable to the human senses. Most common stimulation techniques, such as transcranial magnetic stimulation (TMS) and transcranial alternating current stimulation (tACS) use electromagnetic stimulation to induce electrical currents in brain tissue or to influence resting membrane potentials [1, 2]. Low-intensity focused ultrasound pulsation (LIFUP) stimulates neural tissue using acoustic energy, but at auditory frequencies which are imperceptible to humans [3]. However, sensory stimulation which modulates neural circuits beyond the perceptual pathways that are trivially activated by visual, auditory, or other sensory stimuli may represent a frontier direction in noninvasive brain stimulation [4]. For example, periodic visual stimulation shares similar beneficial effects (e.g., enhanced cortical plasticity [5]), risks (e.g., seizure provocation [6]), and relevant parameters (stimulation frequency [7, 8]) with TMS [9].

In some cases, periodic sensory stimulation (PSS) also has novel perceptual effects which are not commonly expected when the same sensory modality is delivered in an ordinary or naturalistic context (i.e., with constant intensity or infrequent pulses). An early example of this phenomenon was first reported over 200 years ago by Purkinje, who waved his out-stretched fingers between the sun and his closed eyelids to create periodic visual stimulation resulting in the perception of geometric patterns and form constants [10, 11]. In fact, such reports by Purkinje and more recently others [8] closely match the geometric form constants that are perceived with closed eyes under the influence of psychedelic or otherwise hallucinogenic substances [7], suggesting that periodic visual stimulation may reveal the elementary organization [12] of visual cortex (e.g., radial symmetric retinotopic structure) in a manner similar to psychedelic drugs like psilocybin [13, 7, 14]. Additionally, 60 Hz visual PSS may even share plasticity-enhancing effects with the dissociative hallucinogen ketamine [15]. In this regard, PSS may straddle the boundary between inducing neural effects that are conventionally achieved using electromagnetic simulation, on the one hand, and pharmacology, on the other hand.

To better understand this novel approach to brain stimulation, we studied periodic audiovisual stimulation (PAVS) in a large sample. While many recent studies have described the visual effects of stroboscopic stimulation [16, 17, 8, 18], the neural effects are comparatively understudied, with just a handful of published reports [19, 20, 21] which have largely focused on examining EEG frequency changes. We thus sought to fill a gap in the literature by rigorously searching for entrainment effects, which have so far only been observed in small, uncontrolled studies [19, 22].

Toward this end, we studied the effects of stimulation in two PAVS conditions with over 80 healthy participants each, one with multimodal synchronization between visual and auditory modalities (henceforth, ‘experimental stimulation’) and another lacking this component (henceforth, ‘sham stimulation’). Both groups were compared to a control condition with no stimulation in which a comparable number of participants were instead instructed to practice focused breathing meditation, which, like the active conditions, may also be considered a non-pharmacological altered state of consciousness [23, 24, 25] that contains trace rhythmic components (i.e. the inherent regularity of respiration). During stimulation, we recorded spontaneous EEG signals to test the hypothesis that neural oscillations are entrained by the frequency of PAVS. Indeed, the findings presented here offer clear evidence of neural entrainment induced by the visual component of PAVS implemented using a stroboscope device (Fig. 1).

**Fig. 1.**
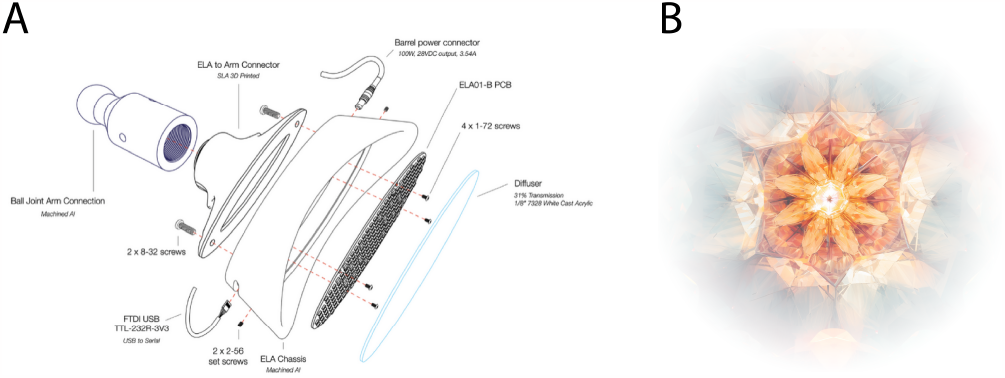
**A) Stroboscopic Device, Exploded View.** The hardware for generating the stroboscopic stimulation was developed by INTO, Inc. and predominantly consisted of a matrix of single-color LEDs that shine through a diffuser that permits 31 percent transmission. **B) Simulated view of a geometric hallucination induced by stroboscopic stimulation.** Image rendered using generative AI software (Midjourney).

## Results

273 adult subjects without photosensitive contraindications completed the experiment. Of these participants, we retained usable EEG data recorded from N = 248 participants. Recordings were analyzed from participants in each of three groups: 1) experimental stimulation (N = 83), 2) sham stimulation (N = 82), and 3) a breath-focused meditation control condition (N = 83). Additionally, participants were assigned to one of three session durations or ‘doses’ independently of the group they were assigned to: 5.5 minutes (N = 81), 11 minutes (N = 83), and 22 minutes (N = 84).

Stimulation frequencies varied throughout the PAVS experience in both experimental stimulation and the sham stimulation groups (Fig. 2). During the experimental stimulation, the frequency of the main strobe (henceforth, ‘strobe A’) varied from 0.14 to 25 Hz, with a median frequency across time of 8 - 9 Hz (8.8 Hz, 5.5-minute session; 8.5 Hz, 11-minute session; 9.6 Hz, 22-minute session). Additionally, a secondary strobe (henceforth, ‘strobe B’) was incorporated which began at 0.3 Hz and then started to flash periodically 1 - 4 minutes into the experience (Fig. 2) with a frequency range of 40.1 to 79.4 Hz; its median frequency across time (excluding the low-frequency floor of 0.3 Hz at the beginning of each session) was 60 - 70 Hz (68.5 Hz, 5.5-minute session; 60.0 Hz, 11-minute session; 59.6 Hz, 22-minute session). Lastly, binaural beats (4.5 Hz - 14.5 Hz) were incorporated with ambient music during each session to add an auditory dimension to the stimulation. The frequency of the binaural beats closely followed the frequency of strobe A (Fig. 2). When present, the median binaural beat frequency across time was 10.5 Hz (5.5-minute session), 8.1 Hz (11-minute session), and 9.3 Hz (22-minute session). Full details can be found in the section labeled “Audiovisual Composition”.

**Fig. 2.**
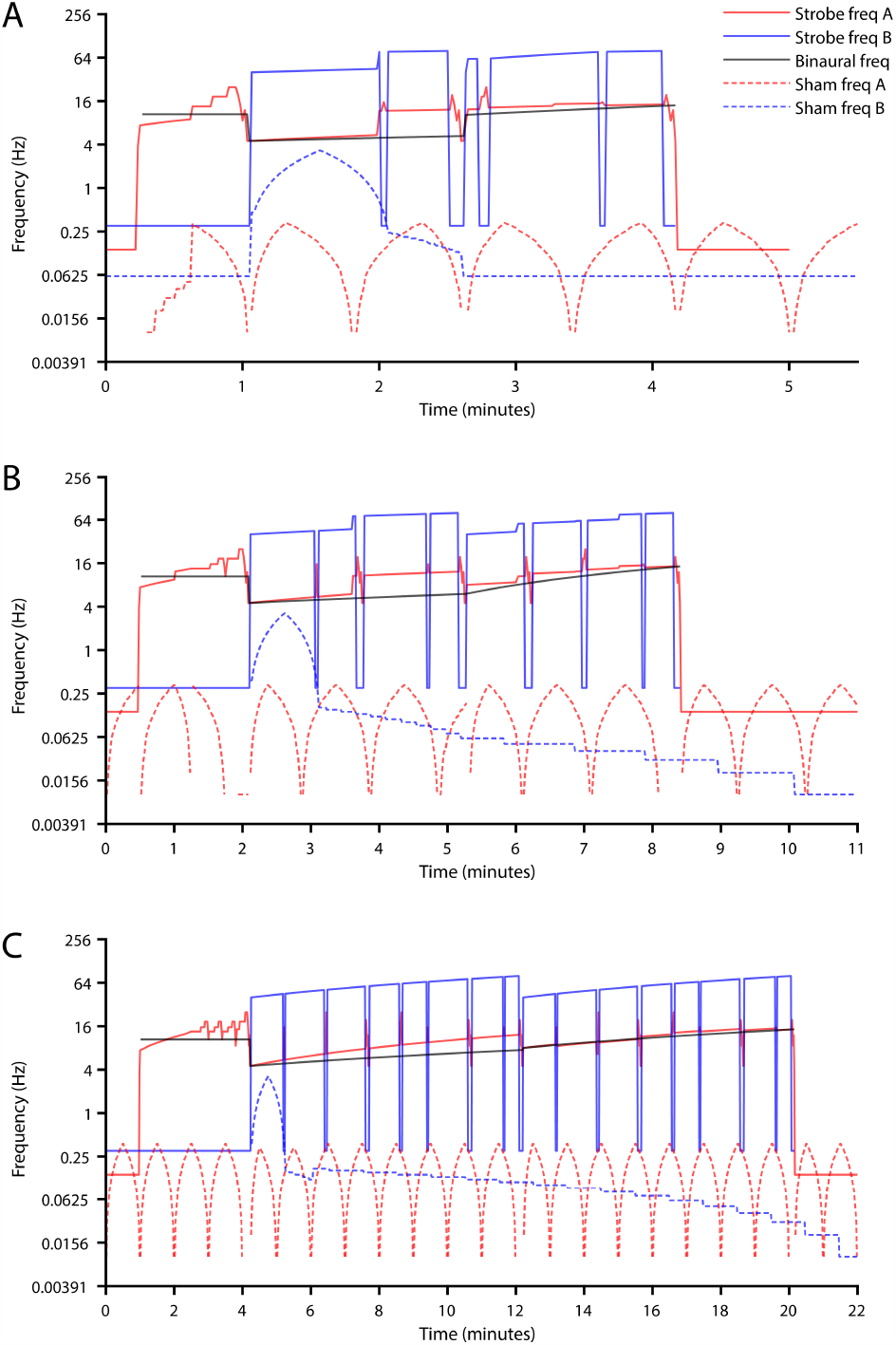
**Stimulation frequencies plotted on a logarithmic frequency scale for different ‘doses’ or session lengths: A) 5.5 minutes, B) 11 minutes, and C) 22 minutes.** The primary strobe frequency (strobe A) is shown in red, while the secondary strobe frequency (strobe B) is shown in blue. The binaural beat frequency (present only in the experimental stimulation condition) is shown in black. Sham stimulation is indicated with a dotted line.

During the sham stimulation, we used much lower stimulation frequencies which were expected to elicit far weaker effects. The frequency of strobe A varied from 0.01 to 0.38 Hz (excluding periods of constant intensity, i.e., 0 Hz stimulation), with a median frequency of 0.15 Hz (5.5-minute session), 0.16 Hz (11-minute session), or 0.19 Hz (22-minute session). Similarly, the frequency of strobe B varied from 0.01 to 3.25 Hz (excluding periods of 0 Hz stimulation), with a median frequency of 0.06 Hz (5.5-minute session), 0.05 Hz (11-minute session), or 1.0 Hz (22-minute session). Binaural beats were absent from the sham stimulation.

We examined the correlation between neural activity and the presence/absence of specific stroboscopic light and audio frequencies, aiming to quantify neural entrainment. The correlation metrics were referenced to the experimental stimulation frequencies, regardless of ‘dose’ assignment. This approach was chosen since the frequencies used in the sham condition were typically below the high-pass filter cutoff frequency (1 Hz), and no stimulation occurred in the meditation control group. Consequently, each comparison group (sham and control) demonstrated the potential level of false positive entrainment that might occur by chance. For a comprehensive description of the correlation computation and the techniques used, please refer to the section labeled “EEG Methods”.

In what follows, all statistics were calculated using an unbalanced (Type III sum of squares), two-way mass-univariate ANOVA following a 3x3 Design: Group + Dose + Group: Dose. We adjusted P-values for testing across multiple channels using false discovery rates applied separately to each ANOVA.

### Stroboscopic stimulation produces widespread neural entrainment

We found a significant main effect of group at all EEG channels for entrainment by strobe A (F > 15, p < 0.001, all channels). Post-hoc tests revealed that this effect was driven by the experimental versus sham contrast (p <0.005, all channels) and the experimental versus control contrast (p < 0.005, all channels). Although this neural entrainment effect is widespread, the largest entrainment effects were detected over visual areas, fitting with the stimulus modality, and lowest over temporoparietal channels (see Fig. 3 and Fig. 4). No posthoc tests were significant for the sham versus control contrast. Dose and Group x Dose interaction effects were not significant for any channels (Fig. 3).

**Fig. 3.**
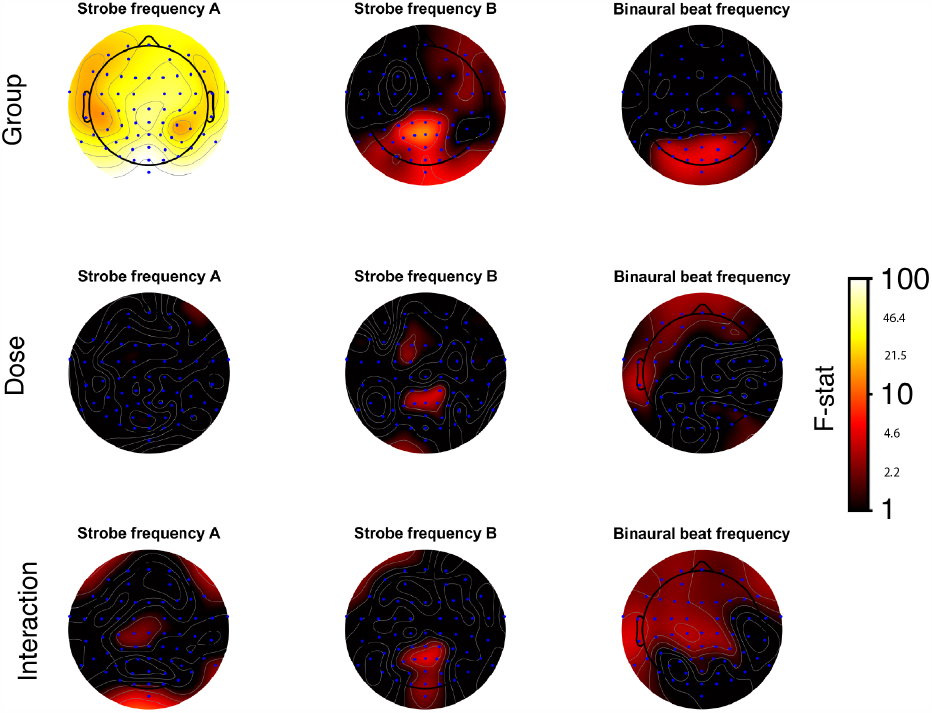
F-statistic ANOVA topographies. Each column is a different contrast and each row considers different stimulation parameters. F-statistics are *log*_10_-scaled given that they span multiple orders of magnitude.

**Fig. 4.**
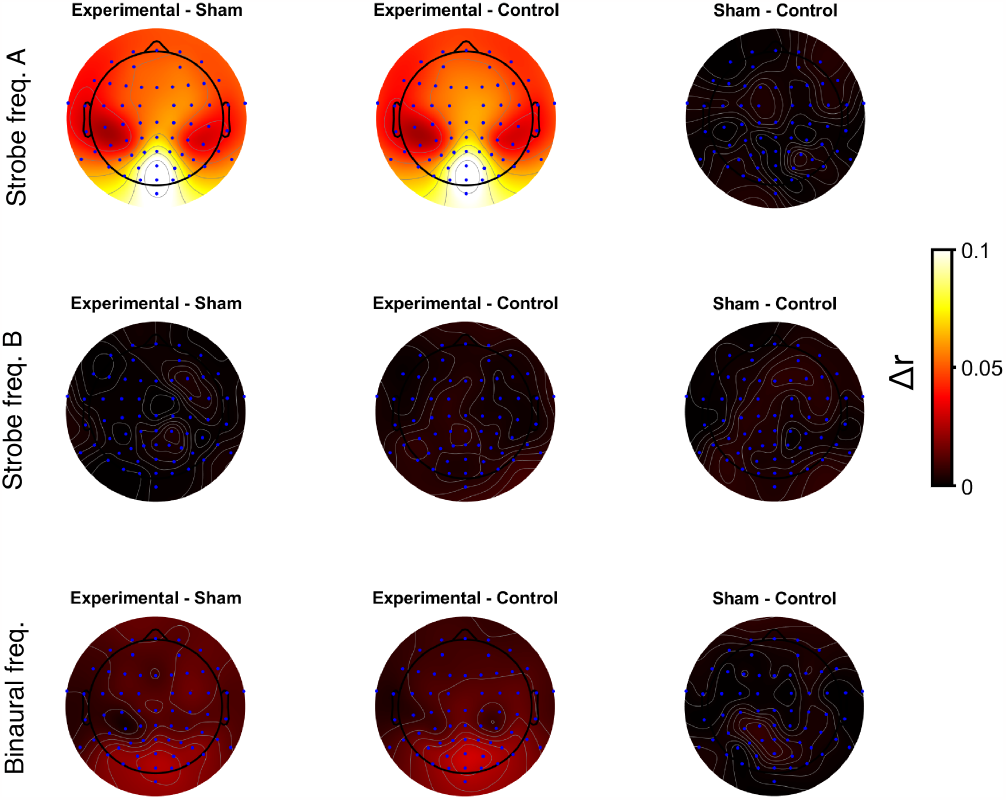
Post-hoc comparisons of correlation coefficients (r). Each column is a different contrast used to compute ∆*r* and each row considers correlations derived using the frequency given by different stimulation parameters.

Although we attempted to measure entrainment effects from strobe B, this proved somewhat impractical as the median strobe frequency (60 - 70 Hz) was above the lowpass filter cutoff frequency (45 Hz). We did not wish to adjust our filter settings, as EEG frequencies above 45 Hz are substantially contaminated by muscle artifacts and electrical line noise in scalp recordings. Nonetheless, we attempted to measure high-frequency entrainment despite these challenges and found several posterior electrodes with a significant main effect of group at the frequency of strobe B (CPz, P3, P2, CP1, POz, Oz, Iz, P < 0.05). This result suggests focal entrainment of posterior areas (e.g., visual cortices). Posthoc tests revealed that three of these channels were significant in the contrast of experimental stimulation with the meditation control (CPz, CP1, Iz, P< 0.05), but no channels were significant in the contrast of experimental stimulation with sham stimulation.

Finally, although we observed what might appear to be evidence of neural entrainment at the binaural beat frequency (Fig. 4), we caution that the binaural beat frequency closely matched the frequency of strobe A (Fig. 2). Given the largely posterior topography of this response (Fig. 4), it is likely that these effects are actually driven by strobe A rather than the binaural beats themselves. Furthermore, we did not observe any statistically significant main effects or interaction with binaural beat frequency (Fig. 3).

### Evidence of a photic driving response

In addition to these quantitative results, we also observed qualitative evidence in EEG waveforms of strong entrainment manifesting as a very well-defined yet benign photic driving response in at least one participant (Fig. 5A), which was independently reviewed by two neurologists. We applied a Morlet wavelet transform to all channels and plotted the channel-averaged spectrogram (Fig. 5B), revealing a burst of spectral energy near the first harmonic of the strobe A frequency, as is common in the photic driving response [26]. Note that because the photic driving response appears non-sinusoidal and was convolved with a Gaussian-windowed sinusoid (i.e., a Morlet wavelet), some spectral leakage and frequency shift may be present, potentially explaining why the response does not occur exactly at 9.2 Hz, i.e, the first harmonic of the strobe A frequency.

**Fig. 5.**
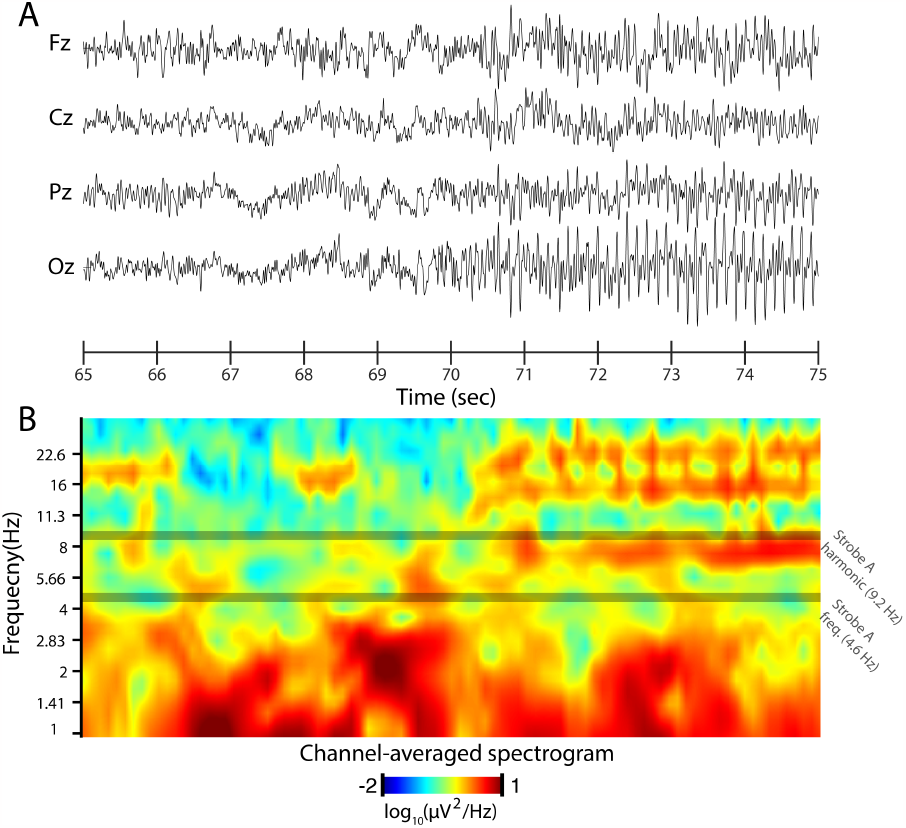
Evidence of photic driving effect. 10 s recordings from 4 midline channels are shown in (A), with the onset of the photic driving effect occurring midway through the displayed time course. We then computed the time-frequency representation of this of these 4 channels using Morlet wavelets and plotted the average time-frequency representation across the 4 channels in (B). The frequency of the main strobe and the first harmonic thereof are noted by dark horizontal lines.

## Discussion

Herein, we have presented strong evidence of neural entrainment caused by the low-frequency (≤ 25 Hz, strobe A) component of PAVS implemented using a stroboscopic device paired with binaural beats (Fig. 1). The entrainment effect appears to occur rapidly, as we did not find an effect of dosage (i.e., session length), and it exhibits an occipitally-dominant topography consistent with the visual nature of the stroboscopic stimulation, but all channels showed significant entrainment effects, likely as a result of volume conduction. We also observed weaker evidence of neural entrainment to higher frequencies (*>* 40 Hz, strobe B) which was limited to parietal and occipital channels. Alongside these entrainment effects, the stroboscopic or “flicker” stimulation utilized in the current study is known to induce geometric perceptions through closed eyes [27, 20, 11, 12], an effect which has received increased attention and research in recent years [16, 7, 5, 8, 17, 18]. Taken together, these findings underscore the potential of PAVS to induce powerful neural effects.

### PAVS as a novel form of noninvasive brain stimulation

Conventionally, noninvasive brain stimulation is implemented using techniques such as TMS or tACS which use magnetic fields (TMS) or electrical currents (tACS) to modulate neuronal excitability [2, 1]. These electromagnetic stimuli, rarely encountered in real-world settings, are generally considered separately from sensory stimuli (including electromagnetic radiation in the visible spectrum), as the brain is nearly constantly exposed to an ambient deluge of these afferent signals during ordinary wakefulness [28]. By comparison, however, PAVS rarely occur in ecological contexts, especially those which were encountered by preindustrial humans. Although humans may self-induce flicker stimulation, e.g., by waving a grated objected or even one’s outstretched fingers between one’s face and a bright light source [10, 11], the phenomenon appears to be sufficiently rare that biological evolution has not prepared the human perceptual system to expect such highly periodic bouts of visual afferent signals, which therefore result in highly non-veridical perceptions, i.e., simple geometric hallucinations similar to those induced by 5*HT*_2*a*_ receptor agonists such as lysergic acid diethylamide (LSD) [29] or psilocybin/psilocyin [7].

These geometric hallucinations might be understood in a predictive processing framework: since the stimuli are not encountered in ecologically valid contexts, the brain lacks a prior model to represent them [30, 14]. However, even after repeated exposure, these geometric perceptions do not disappear, as would be expected in the framework of Bayesian learning, where the brain updates its priors to account for prediction errors. Thus, an attractive lower-level mechanism to explain these geometric hallucinations involves neural entrainment of oscillatory activity in visual cortex which disrupts normal perceptual computations [12]. Support for this hypothesis was obtained from a small EEG study of flicker stimulation over two decades ago which found resonant activity in EEG recorded during the stimulation [19]. However, the study recruited only 10 participants and lacked a control condition. Our work builds on this prior study by increasing the sample size by nearly an order of magnitude, introducing comparison conditions such as sham stimulation and an unstimulated control condition (meditation), and utilizing a more complex, multimodal form of PAVS.

Given our findings, we believe that PAVS can be better conceptualized as a form of noninvasive brain stimulation as opposed to an ordinary sensory stimulus. Like tACS [2], PAVS entrains neural oscillations to the frequency of stimulation. In fact, PAVS reliably and consistently induces geometric phosphenes in participants with little effort on the part of the experimenter [7, 18], whereas visual phosphenes require precise and careful targeting of visual cortex with tACS and TMS [2]. Given that at least one study has suggested that PSS has plasticity-promoting effects [5], we advocate for future work that will compare PAVS (or broader forms of PSS) to TMS as a means of enhancing neural plasticity and potentially treating mood disorders.

### Conclusions, limitations, and future directions

In conclusion, we demonstrated widespread neural entrainment effects, dominant over visual areas, in a sample of over 80 participants stimulated with PAVS as compared with equally-sized samples of sham stimulation and breath-focused meditation. Our results suggest that PAVS has potential as a powerful neurostimulation and/or neuromodulation technology. A shortcoming of our study was that we were unable to also measure possible ultra-low frequency (*<* 0.5 Hz) entrainment relevant to the sham stimulation condition, as the stimulation frequency fell below the frequency cutoff of the high-pass EEG filter. Nonetheless, the sham stimulation condition provided data from which we could estimate the chance level correlation between the experimental stimulation and EEG spectral power. For similar reasons owing to the low-pass EEG filter, we were challenged in our efforts to correlate high-frequency stimulation from strobe B in the experimental stimulation condition with spectral EEG power. Nonetheless, even after applying this necessary filtering, we found effects suggestive of focal entrainment for high frequencies. Finally, we did not collect data on geometric perception during the PAVS experience. However, given that many previous studies [11, 20, 27, 7] have focused on the perceptual aspect of stroboscopic stimulation, we chose to focus instead on neural effects which have been barely explored until the present.

Based on our results, we advocate for future studies that will directly compare PAVS to established forms of neurostimulation such as TMS as a means of entraining neural activity, enhancing neural plasticity in cortical circuits, and perhaps even treating mood disorders such as major depression.

## Materials and Methods

### Experimental Methods

#### Participants

286 participants were recruited for this study by way of Facebook ads targeting adults within a 50-mile radius of Santa Monica, California, USA. 13 participants were either excluded or unable to finish the study in its entirety due to voluntary withdrawal from the study, technical difficulties, or falling asleep. As such a total of 273 participants, aged 19-79 (M = 43.73; SD = 15.58; 142 females) completed this study. Participants were compensated at a rate of 30 US dollars per hour via cash or the mobile payment service Venmo, and the parking fee was waived for all participants.

Participants were not permitted to participate in this study if they had a history of epilepsy and/or seizures, migraines, photo-light sensitivity, cataracts, corneal abrasions, keratitis, uveitis, hearing problems, or non-normal/non-corrected vision. Participants were also excluded if they were currently taking any photophobia-inducing medications or hearing-altering medications. Eligibility screening was conducted prior to the participant’s enrollment in the study using Castor ePRO (Amsterdam, Netherlands). All participants digitally signed an informed consent using Castor eConsent. The Advarra (Columbia, Maryland, USA) Institutional Review Board approved all recruitment and testing procedures prior to initiating enrollment (Pro00048382).

#### Materials

EEG signals (500Hz sampling rate) were acquired using a dual-amp 64-channel cap system (BrainVision, LLC, Santa Fe, NM) connected to a 15.6” 2021 Lenovo Ideapad. Measurements of the participant’s head from their nasion to inion determined which cap size (54cm, 56cm, 58cm, or 60cm) was utilized. The cap was positioned on the participant’s head such that channel FPz was at 10% of the distance from nasion to inion, midline channels were aligned, and the velcro chin strap was taught, but comfortable. Nuprep skin prep gel (Weaver and Co.) was used to exfoliate the scalp through electrodes before applying Neurospec abrasive electrolyte gel (EasyCap, Inc.). All powered devices, with the exception of the stroboscopic device, were unplugged prior to the experiential portion of the experiment to help prevent the impact of line noise on the EEG data.

The stroboscopic device (INTO, Inc., Santa Fe, NM) was an initial prototype of a phosphene generation device that uses an array of 8 color frequencies among 192 LEDs to output light through a 31% opacity diffuser (Fig. 1). The LEDs were programmed to pulse at specific frequencies and the dynamic, time-varying patterns were paired with a pre-recorded stereo audio track. The combination of the LED pattern and audio tracks is termed an experience herein and the exact compositions can be found in the Audiovisual Composition section below.

Wired earbuds (Sony XBA-100) were provided to the participants to place in their ears. The device was positioned 12.7 cm (5 inches) away from the participant’s nose using a desk mount swivel (M!ka). All participants sat in a powered recliner chair regardless of group assignment and were instructed to adjust the leg and back positions to their comfort. All window shades were lowered prior to the start of each experiential portion of the experiment. Participants were instructed to keep their eyes closed throughout the duration of the light stimulation as the experience was intended to output light onto the eyelids (see Fig. 6).

**Fig. 6.**
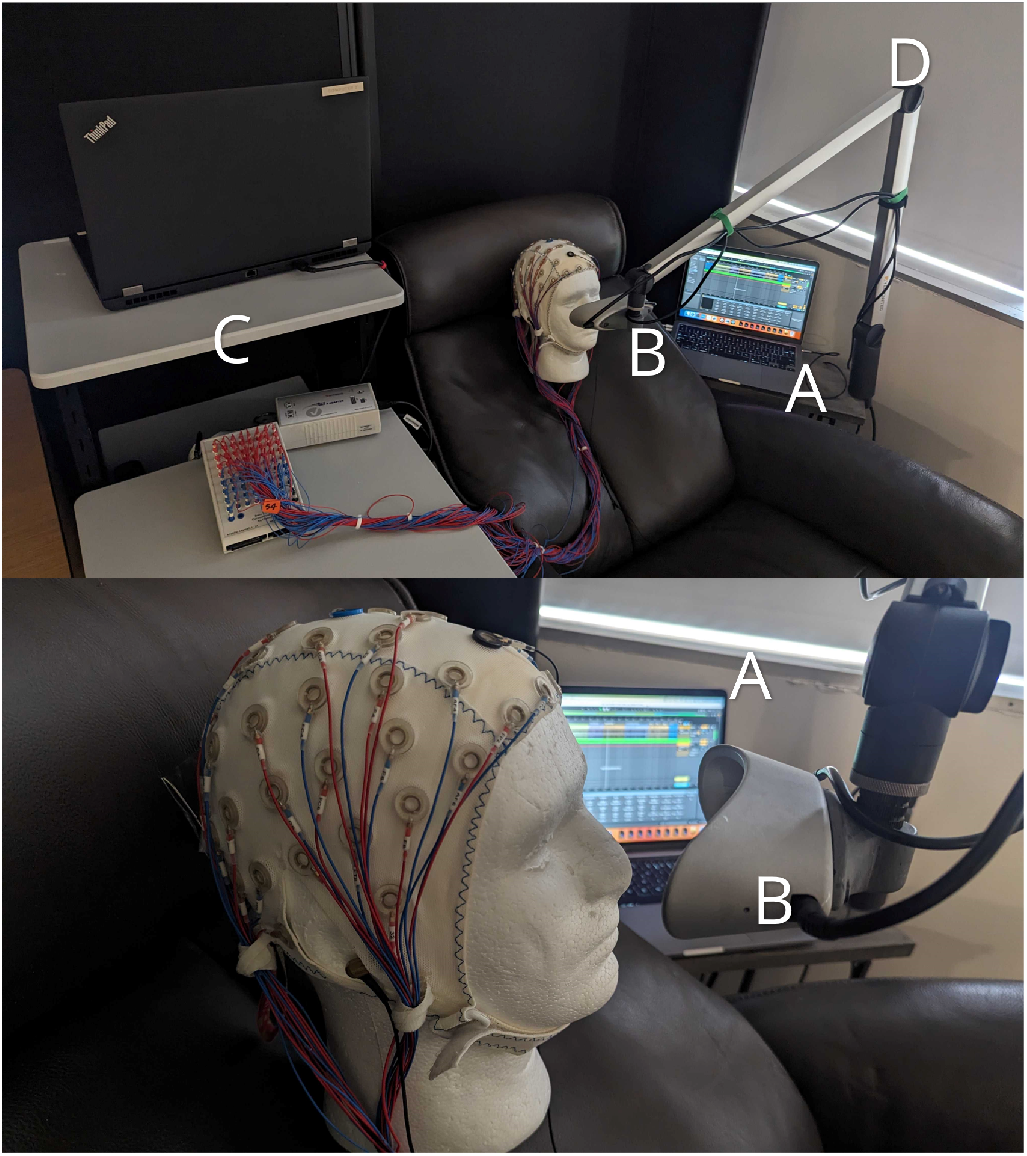
Experimental setup (stroboscopic conditions) Top Panel: (A) Macbook Air running the Ableton 10 audiovisual experience controller (B) Stroboscopic device (C) 64-channel EEG computer system (D) M!ka extension arm to hold the device. Bottom panel: a side view of A and B positioned 12.7 cm from the participant’s closed eyes.

Experiential compositions were triggered using Ableton Live 10 via a Python3 Controller, Pylive (https://github.com/ideoforms/pylive) on a 13.3” 2020 Macbook Air. Lab Streaming Layer with LabRecorder (https://github.com/labstreaminglayer) was utilized to temporally synchronize our EEG, peripheral, and experimental time series (e.g. pre-experience rest ended, Ableton experience started) within an XDF file format.

Randomization was conducted using a single-site validated block randomization model (Castor EDC) with gender as a randomization strata across 9 groups (3 experiences x 3 duration periods). Both the participants and experimenters were blind as to group assignment until after the first resting state period.

#### Procedure

Upon arrival, all participants were temperature screened using an infrared no-touch thermometer (iHealth Labs Inc.) and offered an N-95 face mask if they arrived without a mask of their own. Participants were then seated on an office chair facing a 90.2 cm x 127 cm desk and given an overview of the experimental session (i.e. outfitting of EEG and peripherals, rest, “experience”, rest) and told their “experience” would require them to sit for a period of 5.5, 11, or 22 minutes as they either did a breath-focused meditation exercise or received light and sound stimulation. Neither the participant nor the experimenter was aware of which experiential group the participant would be assigned to at this time.

Participants were instructed to put their phones on silent, remove all jewelry, and remove all bulky items from their person to maximize their comfort. Participants were outfitted with the EEG cap, bio-peripherals, and earbuds (regardless of group) while seated in a recliner chair. Participants were then instructed to close their eyes and relax, trying their best not to fall asleep, for 5 minutes. Afterward, the experimental script would reveal the randomized group assignment to the participant and experimenter for the first time. Participants were assigned to either Audiovisual Experience 1 (Experimental), Audiovisual Experience 2 (Sham), or a Control group (breath-focused meditation) with a sub-group of either 5.5, 11, or 22 minutes, for a total of 9 groups.

If assigned to an audiovisual experience, participants had the device positioned in front of their closed eyes (see materials above) before the experimental script triggered the launch of the experience. Subjects were told they could easily swivel the mounting arm and exit the experience at any time if they so desired. If assigned to the meditation group, participants were read instructions for a breath-focused awareness meditation before engaging in the meditation in the same seated position as the other groups. Following this experiential period, participants were able to open their eyes briefly before engaging in a second period of 5-minute closed-eye rest. Afterwards, the EEG and bio peripherals were removed and participants were permitted to use the restroom to rinse their hair. Following this cleanup, participants completed a post-experience behavioral assay before being compensated and dismissed from the study.

### EEG Methods

All analyses were focused on the spontaneous EEG data collected throughout the duration of the PVAS.

Exclusions were assessed based on testing protocol abnormalities provided by INTO and further outliers were identified from signal quality, missing data, or frequency band power statistics. All subjects identified as outliers were removed from group statistics and the number of subjects in each group for each phase of the analyses is listed in Table 1 below.

**Table 1.** Sample Sizes Stratified by Group Assignment and Intervention Duration.

|  | Group | # of Participants per Group |
| --- | --- | --- |
| 5.5 minutes | Experimental | 25 |
|  | Sham | 28 |
|  | Control (Meditation) | 28 |
| 11 minutes | Experimental | 30 |
|  | Sham | 25 |
|  | Control (Meditation) | 28 |
| 22 minutes | Experimental | 28 |
|  | Sham | 29 |
|  | Control (Meditation) | 27 |

#### Pre-Processing

The data were subjected to a standard preprocessing chain in NeuroPype™ which removes and/or repairs data from corrupted electrodes and signal artifacts (drift, line noise, and high-variance artifacts including blinks, movement, cardiac, and muscle) and prepares the data for subsequent analysis. This includes the following stages: 1) Inferring channel locations from their 10-20 labels; 2) Removal of channels that have no location; 3) FIR high-pass filter with a transition band between 0.25 and 0.5 Hz; 4) Removal of bad channels using a correlation and high-frequency noise criterion similar to the PREP methods [31]; 5) Removal of high-amplitude artifacts using Artifact Subspace Reconstruction [32] with an artifact threshold of 20 standard deviations; 6) Removal of residual high-artifact time windows using a variance criterion; 7) Electrode re-referencing to Common Average Reference; 8) Independent Component Analysis (ICA) using FastICA [33] ; 9) Automated classification of and rejection of bad components (eye, muscle, line noise, cardiac) using ICLabel [34]; 10) Spherical-spline interpolation of removed channels; 11) FIR low-pass filter with a transition band between 45 and 50 Hz.

#### EEG Spectral Analysis

EEG spectrum correlations with the stroboscopic light and audio frequencies were computed for each EEG channel. EEG power spectrum was computed with a multitaper method and utilized a four-second epoch window (with a one-second sliding window) to establish one-second epochs with 1 Hz frequency bins without overlap into adjacent frequencies.

#### Experiential Composition Correlations (Entrainment)

To quantify neural entrainment, we correlated the binary presence/absence of the strobes and audio at a given frequency taken from the experimental stimulation condition with EEG spectral power at the corresponding frequency. In order to match the same EEG spectrum 1 Hz bins, the stroboscopic light/audio frequencies were rounded to the nearest 1 Hz. For each 1 Hz bin, we created a square wave over time (at one-second intervals) aligned to the EEG spectral time series to indicate when a given frequency was presented (high) to the participant (separately for each stroboscopic light/audio source) and set to 0 (low) when it was not present. Each square wave for each frequency bin was then correlated with the corresponding EEG frequency bin time series across the “experience” to compute a Pearson r correlation coefficient for every 1 Hz bin. Notably, since the sham stimulation and control condition groups did not share the same stroboscopic light/audio frequencies and timings as the experimental stimulation group, we used the same generated square waves over time from the experimental stimulation group (matching the correct dosage group) and performed the same correlation procedure for the sham stimulation and control condition groups. For each EEG channel, we averaged the r-values across all frequency bins for each of the 3 stroboscopic/audio sources (strobe A, strobe B, and audio frequencies). This process resulted in each channel having a single mean r-value per stroboscopic/audio source for the entire session. Group statistics and post-hoc tests used these session mean r-values per channel by stroboscopic/audio source.

#### Photic driving response

We analyzed the photic driving response in Fig. 5 using Morlet wavelets to generate a time-frequency representation of the signal. We used 41 logarithmically spaced wavelets (8 per octave) ranging from 1.0 to 32 Hz, inclusive.

#### Statistical Analyses

Entrainment analyses used the entire “experience” condition for each of the EEG metrics (e.g. Pearson correlation coefficients). Group-level statistics analyzed the effects of Group and Dose factors using the following 3-level full factorial design:

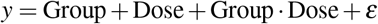

This design was computed with an unbalanced (Type III sum of squares), two-way mass-univariate ANOVA. We corrected for multiple comparisons (False Discovery Rate [35]), and this was done separately for a priori and post-hoc tests. Post-hoc pairwise comparisons used Tukey HSD to account for familywise error.

### AudioVisual Composition

In both the experimental and sham conditions, the same musical composition was used, effectively controlling for potential cognitive and affective confounds that might have been introduced with the use of different auditory stimuli. The music was in the key of D, with a D 11th chord. The binaural beat carrier frequency followed the key changes D and B minor. The musical tone D was chosen for its use in various cultural and spiritual traditions. Using D as the key center allowed the audio composer (Jeff Bova, INTO, Inc.) to “provide a cognitive, psychological reference known for its use in meditation and therapeutic uses, which promoted relaxation, meditation, introspection, and other positive attributes”. This permitted the subject to attune gently to the key of D before entrainment commenced. Furthermore, the musical piece ended with a return to the warmup tonality of the D 11th Chord, with the intention to permit the subject to return to a more wakeful consciousness while integrating their experience.

The primary auditory distinction between the two compositions lay in the presence (experimental) or absence (sham) of binaural beats intended to induce auditory entrainment, with the intent to ensure that any differences in subjects’ responses were attributed purely to entrainment effects and not to individual biases towards the musical composition itself. As for the visual component, both conditions presented similarly to naive observers. However, the rhythmic light patterns differed as a function of composition based on their hypothesized potential to induce entrainment; the sham light maintained a slow and consistent pattern, moving between 0.2 Hz and 0.22 Hz, a range that the composers suspected would not promote light-based entrainment. Importantly, the total lumen output over the course of the experiences was matched, with the intention to isolate the variable of entrainment in subsequent analyses. More details about each composition are included below.

#### Experimental Composition

Briefly, the experimental condition was designed with the intent to encourage a state of relaxation and included the auditory composition described above, a light composition, and binaural beats. The visual composer, Stephen Auger (INTO, Inc.), relayed that the selection of the light flickering frequencies used in the experimental condition was inspired by the subtle pulsations of flames caused by pressure waves of shamanic drumming– an auditory experience known in and of itself to induce trance-like meditative states [36] Table 2 outlines in detail each stage of the experimental composition across all three length variations. The frequency band nomenclature utilized below was borrowed from human brainwave frequency ranges, given the explicit intent to entrain neural patterns of activity: theta (4-8Hz), alpha (8-13Hz), beta (13-32Hz), gamma (32-100Hz).

**Table 2.**
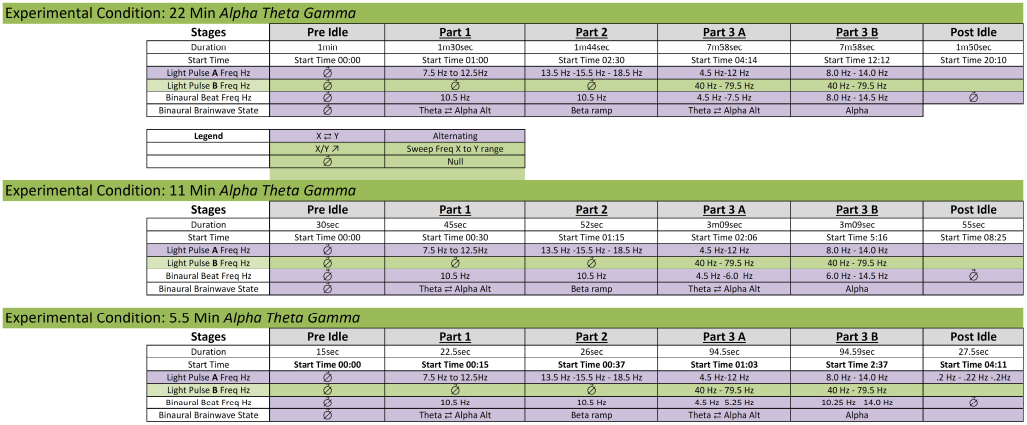
Audiovisual Composition: Experimental Condition displaying each epoch (part) of the experience with duration, start times, light pulse frequencies for both A and B pulses, and binaural beat frequencies with the intended brainwave outcome.

First, in a “pre Idle” section, the visual experience was initiated with slow wave-like patterns that allowed the subject to attune gently to the sensation of light through their closed eyes before the entrainment period commenced, initially pulsing at > .4 Hz speeds, chosen to theoretically induce relaxation and introspection.

Subsequent “entrainment” sections included four stages:

1. Part 1. Light patterns flickering alternating within the theta and alpha frequency ranges.
2. Part 2. The light modulation becomes complex, using increasingly higher beta frequencies.
3. Part 3A. The auditory tone shifts, intending to reflect a deeper, reflective feeling. The light frequencies oscillate across theta and alpha. A secondary light pattern also emerges, oscillating within gamma frequencies.
4. Part 3B. In the final stage, light patterns oscillate across theta, alpha, and beta. In tandem, a second pattern oscillates within gamma frequencies.

At the end of the piece, the ‘post idle’ goes back to the slow, wave-like patterns of the ‘pre idle’. This matches the music’s goal of guiding listeners back to a more alert state.

### Sham Composition

The sham condition is outlined in Table 3. and features an asynchronous series of pulsing light frequencies that modulate the light frequencies at irregular intervals, oscillating between 0.2Hz and 0.22Hz– a frequency range chosen with the intent to not facilitate light-based entrainment. The visual composer relayed that the consistent and non-varying frequency was utilized in an attempt to ensure that any cognitive or emotional reactions elicited by the light were not due to entrainment but merely the presence of light itself.

**Table 3.**
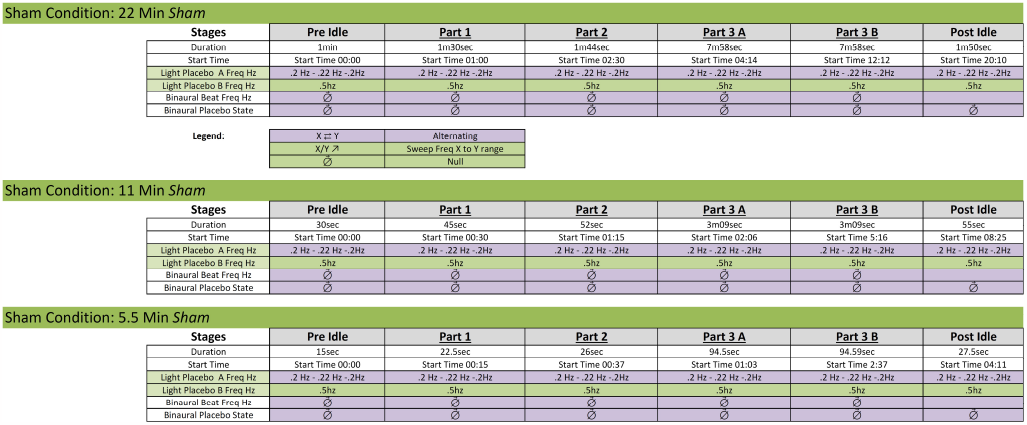
Audiovisual Composition: Sham Condition displaying each epoch (part) of the experience with duration, start times, light pulse frequencies for both A and B pulses, and binaural beat frequencies with the intended brainwave outcome. Note the absence of binaural beat frequencies in the sham condition.

## Acknowledgments

We warmly thank our research participants. We are also grateful to Inna Keselman and Mark Nespeca for reviewing EEG from a photic driving response. We also relay thanks to the team of research assistants at IACS that assisted with data collection: Amir Vala Tavakoli, Jimmy Cenci, Geena Wang, and Caitlin Lynch. This work was funded by a Research Services Agreement between IACS and INTO, Inc.. INTO had no input on the design nor analysis of this study, apart from supplying the stroboscopic hardware and software.

## Author contributions statement

Study Conceptualization and Design: N.R. was responsible for the initial conceptualization and design of the study. Study Implementation: N.R. and N.S. collaborated on the implementation of the study, including data collection and experimental setup. EEG Analyses: J.F., G.H., and C.K. conducted the EEG analyses, interpreting the data and contributing to the analytical framework. Manuscript Writing (1st Draft): N.R. and J.F. co-wrote the first draft of the manuscript, integrating the study design, implementation, and initial findings. Manuscript Writing (Review and Edits): The manuscript was reviewed and edited by J.F., N.R., N.S., G.H., and C.K. All authors contributed to the refinement of the manuscript and approved the final version for submission.

